# DeepMQ: A Deep Learning Approach Based Myelin Quantification in Microscopic Fluorescence Images

**DOI:** 10.1101/407643

**Authors:** Sibel Çimen, Abdulkerim Çapar, Dursun Ali Ekinci, Umut Engin Ayten, Bilal Ersen Kerman, Behçet Uğur Töreyin

## Abstract

Oligodendrocytes wrap around the axons and form the myelin. Myelin facilitates rapid neural signal transmission. Any damage to myelin disrupts neuronal communication leading to neurological diseases such as multiple sclerosis (MS). There is no cure for MS. This is, in part, due to lack of an efficient method for myelin quantification during drug screening. In this study, an image analysis based myelin sheath detection method, DeepMQ, is developed. The method consists of a feature extraction step followed by a deep learning based binary classification module. The images, which were acquired on a confocal microscope contain three channels and multiple z-sections. Each channel represents either oligodendroyctes, neurons, or nuclei. During feature extraction, 26-neighbours of each voxel is mapped onto a 2D feature image. This image is, then, fed to the deep learning classifier, in order to detect myelin. Results indicate that 93.38% accuracy is achieved in a set of fluorescence microscope images of mouse stem cell-derived oligodendroyctes and neurons. To the best of authors’ knowledge, this is the first study utilizing image analysis along with machine learning techniques to quantify myelination.

## I. INTRODUCTION

Generally, increases in our understanding of the nature follow advances in technology. For example, Leuwenhoeks invention of the microscope brought light on to an invisible world. At present, advances in microscopy enabled relatively easy acquisition of images raising a new limitation: extraction of meaningful data from the images [1]. When the time and man power required for image analysis exceeds available resources, life scientists turn to automation. Recently, in a related work, a team of researchers at the Allen Brain Institute reconstructed neurons from a large volume of brain images [2]. On the other hand, myelin, another structure within the nervous system, poses a different question, because it is composed of parts from two different cell types: the axon of the neuron and the process of the oligodendrocyte (cf. Fig.1). Life scientists identify myelin via the shape and the configuration of overlap between these two structures [3]. An experienced researcher can annotate myelin over different imaging planes and can distinguish it from non-myelin overlaps between cells protrusions (cf. Fig.1). For a large number of images or images covering a large area, this process can take hours to days [3]. Thus, automation of myelin quantification will not only save a large amount of researcher hours, but is also a challenging image analysis question.

**Fig. 1.**
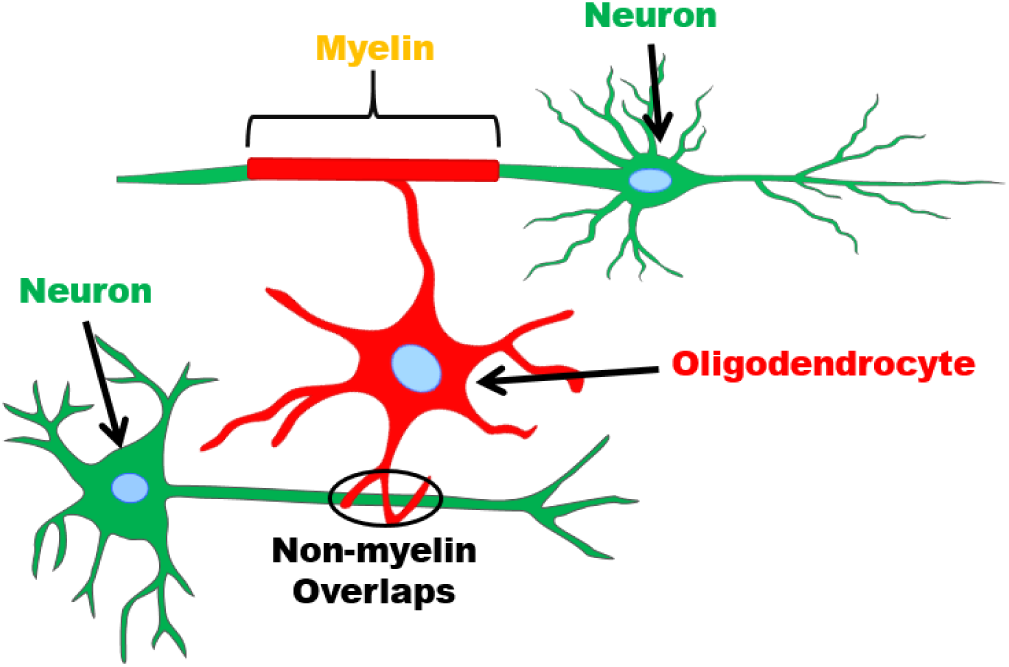
Demonstration of myelin and non-myelin. Oligodendrocytes wrap around axons to form myelin sheaths. However, not all the overlap between oligodendrocytes and the neurons correspond to myelin sheaths.

In many cases, the methods to analyze the microscopic co-localization are often simple and descriptive [4]. The overlap of the two signals are assessed on false colored images. Such a visual evaluation requires comparable fluorescence intensities of the two markers, may be subject to biases, and is hardly quantitative. Recognition of these issues quite early in the development of fluorescence microscopy led to use of statistical parameters to evaluate the correlation of fluorescence-intensities of two (or more) detection channels on a pixel-by-pixel basis [4]. For the confocal microscopy, a method [9] was presented to visualize the total overlap of two patterns by subtracting one component of an image from the other and this method provides a clear representation of complete overlap. However, simply identifying co-localization is inadequate to segment myelin because the cell protrusions may overlap resulting in false positives (cf. Fig.1) [3]. Thus, we employed a machine learning-based approach in order to automate myelin quantification. To that extent, SVM, DT, and deep learning based methods were utilized on feature images. Feature images are obtained by expressing three dimensional spatial relation between neighboring voxels in two dimensions. To the best of authors’ knowledge, this is the first study that aims at quantifying myelination using image analysis and machine learning methods. Having said that, there are studies on three-dimensional medical image segmentation and quantification [5]–[8].

The paper is organized as follows. Myelin and the need for automated quantification is discussed in Section II. Section III describes Materials and Methods while Section IV describes Classification Methods. Experimental Results are presented in Section V. Conclusion is discussed in Section VI.

## II. MYELIN AND THE NEED FOR AUTOMATED QUANTIFICATION

In the vertebrate nervous system protrusions of the oligodens drocytes wraps around the protrusions of the nerve cells, the axons, forming the myelin (cf. Fig.1). The insulation provided by the myelin increases speed and efficiency of the signal transmission across the neurons and supports the survival of the neurons [10]. Myelin is a vital structure for the function of the nervous system, thus, any damage to myelin disrupts the life of the individual leading to neurological diseases, such as, multiple sclerosis (MS). MS is a neuroinflammatory disease of the central neurvous system CNS affecting approximately 2.5 million individuals worldwide [11], [12]. Body’s own immune system attack and destroy myelin [12], [13]. Current therapies reduce demyelination by suppressing the immune system but do not enhance remyelination and, thus, fail to cure the disease [11], [12], [14]. The innate myelin regeneration of the nervous system is inadequate to overcome the destructive potential of the immune attack [9]. In order for the nervous system to regain its function, identification of novel drugs that modulate the immune system and promote rebuilding of the myelin are required. The drug development process usually starts with testing thousands or hundreds of thousands of compounds in a disease relevant assay [15]. However, as myelin quantification is time consuming and labor intensive, it is not feasible to screen for such a high number of compounds. Therefore, rapid quantification of myelin will expedite drug development for MS and other myelin diseases by enabling testing of a large number of candidate chemicals quickly [11]. Additionally, the newly available automated tracing of the neurons from ultra-large images will assist in describing the synaptic connections in the nervous system [2]. When employed on time lapse images this tool may even highlight the changes in neuronal connectivity over time. However, describing the neuronal pattern is only half of the story as myelin influence neuron function [10]. Changes in myelin pattern in the brain is part of the plasticity of brain in response to learning and life experiences [16]. Detecting and cataloguing changes in myelin along with the neurons is critical in understanding the function and plasticity of the nervous system. Thus, an automated myelin quantification method is essential to complement the advances in automated reconstruction of neurons.

## III. MATERIALS AND METHODS

The data used for this study are published in [3]. Briefly, mouse stem cell-derived oligodendrocytes and axons were grown in microfluidic device co-cultures optimized for myelination. Images of an entire chamber of the microfluidic device were acquired on a Zeiss LSM confocal microscope. A sample image is shown in (cf. Fig.2).

**Fig. 2.**
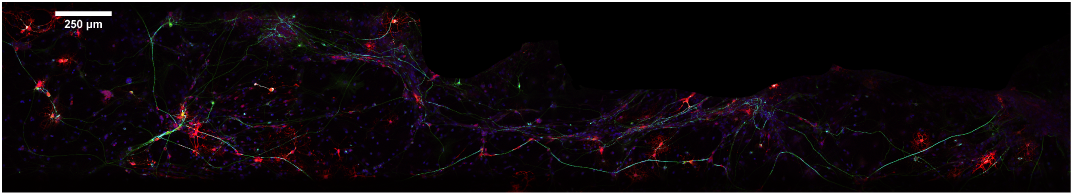
A representative image of a myelinating co-culture. In this maximum intensity projection of the microscopic fluorescence multi-channel image, oligodendrocytes are in the red channel, axons are in the green channel and blue channel is for the nuclei. Scale bar: 250*μ*m

In order to accelerate myelin quantification, one of the authors of the current study, previously, developed the Computer-assisted Evaluation of Myelin (CEM) software [3]. CEM performed operations on fluorescence images and was implemented mainly on the ImageJ platform [17] to be available to a wide range of users. It was able to identify and quantify myelin from large data sets and to detect changes in myelin formation [3], [18]. The algorithm for identification of myelin formation was based on detection of co-localization between neurons and oligodendrocytes. To perform calculations, the images were first converted to binary images. The optimal threshold was determined manually by the researchers. Binary images were further processed to remove cell bodies of oligodendrocytes and neurons. The overlap between the resulting images was identified as ‘myelin’ (cf. Fig.3). Despite the removal of the cell bodies, some overlap between neurons and oligodendrocytes remained resulting in false positive identification (cf. Fig.3). Finally, to calculate the total amount of ‘myelin’, overlapping pixels were counted (cf. Fig.3) [3]. In the current study, myelin identified by CEM was curated by a researcher and was used as the gold standard.

**Fig. 3.**
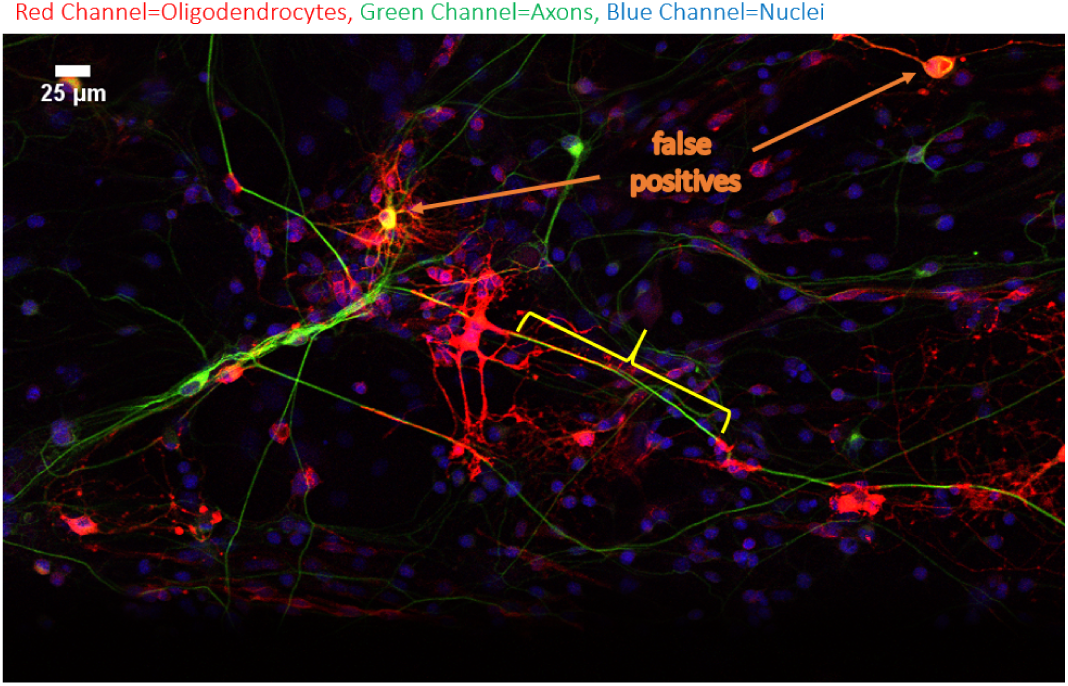
A closeup view of the boxed region in (cf. Fig. 2). Oligodendrocytes (Red Channel), Axons (Green Channel), and Nuclei (Blue Channel) can be identified. Bracket shows a myelin region while the blue lines point to non-myelin overlaps i.e. false positives. Scale bar: 25*μ*m

**Fig. 4.**
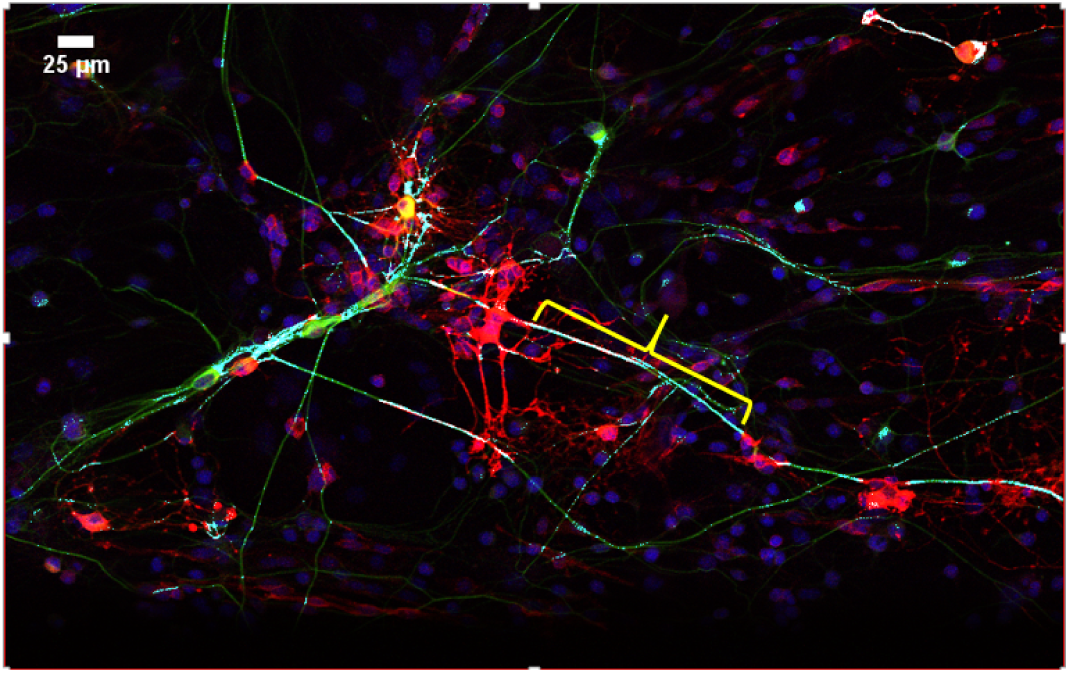
CEM identified the myelin (bracket) in a magnified view of the (cf. Fig. 3). The overlaping regions (Cyan Channel) between oligodendrocytes (Red Channel) and axons (Green Channel) that were segmented by CEM. Nuclei (Blue Channel) are also visible. Scale bar: 25*μ*m

As described above, myelin quantification by CEM is a very simple pixel counting process, which does not take the 3D shape of the myelin into account in identification of myelin regions. Despite the best efforts to minimize the false positives, some still exist (cf. Fig. 3). Additionally, CEM calculates neither the length, nor the number of myelin segments. In the current study, we aimed to overcome these shortcomings by utilising a machine learning-based approach.

First, 3×3 matrices were formed for each channel (red, green, blue) carrying the information for the oligodendrocytes, neurons and nuclei (cf. Fig.5). In order to find myelin regions in 3D, z-stack continuity is considered by forming 3×3 matrices for the immediate z-section neighbors, namely, (z-1) and (z+1). These matrices are merged together and reduced to 2 dimensions, resulting in a 9×9 image (cf. Fig.5) that is expressed as (1). We termed these feature images ‘spectro-spatial feature images’.

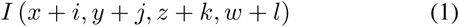

**Fig. 5.**
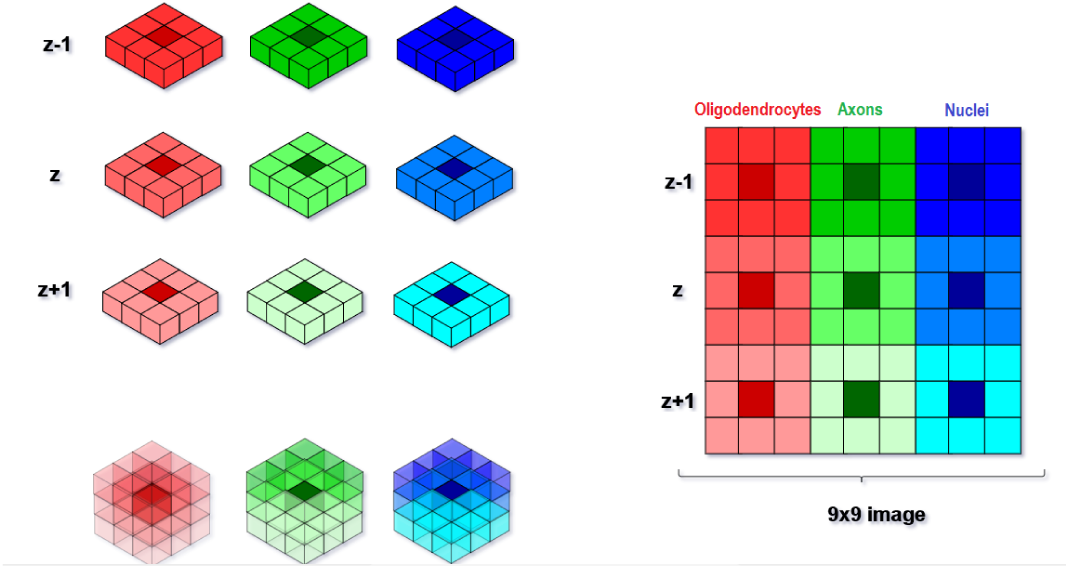
Feature image, corresponding to each voxel, is composed of oligodendrocytes (red), axons (green), and nuclei (blue) channel intensity values of the immediate 26-neighbor voxels.

In the feature images, myelin images are classified as positive, while non-myelin images are classified as negative. The number of feature images obtained is 5364 negative, 5404 positive. Negative and positive samples are grouped into training and test data sets. Details of the data for training and testing are shown in Table I.

**TABLE I.** NUMBER OF DATA

|  | Positive | Negative |
| --- | --- | --- |
| Feature Vectors | 5404 | 5364 |
| For training | 2702 | 2682 |
| For testing | 2702 | 2682 |

Next, the 9× 9 feature images are magnified digitally to 27× 27 in order to increase the performance and to be more effective. The portion of the image to be magnified is selected and then each digital sample comprising the portion to be magnified is repeated in both the horizontal and the vertical direction to expand the selected portion of the image [19]. Each pixel of the original feature image patch is repeated two more times in both the horizontal and the vertical direction (digital magnification). This way the feature image of size 9×9 pixels is expanded to 27×27 pixels. These feature images are shown in (cf. Fig.6).

**Fig. 6.**
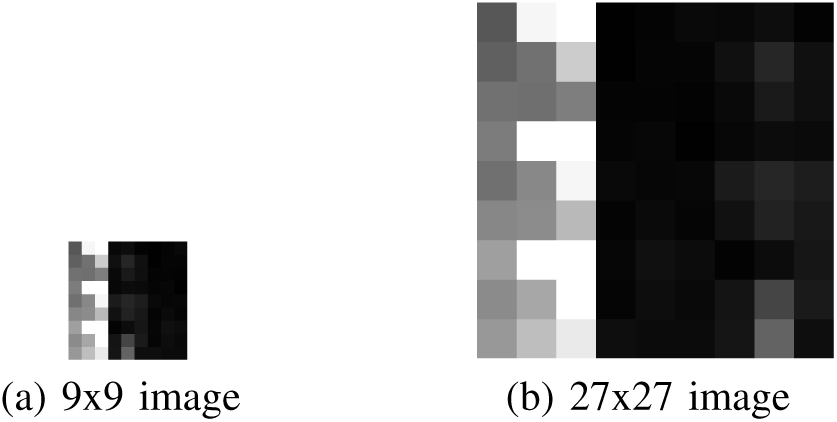
Feature images. a) 9×9 feature image is created from 3 channels (R,G,B). Each channel has 3× 3× 3 images, which is reduced to 2 dimensions.b) 27× 27 feature image is obtained from 9× 9 image by digital magnification method.

## IV. CLASSIFICATION METHODS

There are a plenty of efficient classification methods that one may benefit from in the machine learning literature. Due to their relatively better performances in various fields of research, Support Vector Machines (SVMs), Decision Tree (DT) and Deep Learning (LeNet) are utilised for myelin classification.

One of the most successful machine learning algorithms developed for solving classification problems in recent years is Support Vector Machines. SVMs have been successfully applied to solve many classification problems and have taken place in the literature as one of the efficient machine learning algorithms with high generalization performance [21]. The main advantage of SVMs is to convert the classification problem into a square optimization problem and solve it. Thus, the number of transactions decreases during the learning phase related to the solution of the problem and SVM achieves faster solution than other techniques / algorithms. In addition, optimization-based point classification performance, computational flexibility and adaptability are successful [21].

DTs are also supervised learning method used for classification and regression. The goal is to create a model that predicts the value of a target variable by learning simple decision rules inferred from the data features [22]. Decision trees are a form of tree structure that can be built on both regression and classification models. Classification is used on categorical data such as yes/no while regression is used on numeric target data.

Convolutional Neural Networks (CNNs) are special kind of multi-layer neural networks. They are trained with a version of the back-propagation algorithm where they differ is in the architecture. To recognize the visual patterns from the pixels with minimum pre-processing, CNNs are designed. They can recognize patterns, and with robustness distortions and simple geometric transformations [20]. In this paper, we used the LeNet network, which is known to work well on digit classification tasks. LeNet-5 is a CNN, which was designed for handwritten and machine-printed character recognition. LeNet was described in Gradient-based learning applied to document recognition. LeNet-5 is a CNN, which works on 28×28×1 input data [20]. Also, Caffe is the open framework, which is used in LeNet. Expressive architecture encourages application and innovation. Models and optimization are defined by configuration without hard-coding. Switch between CPU and GPU by setting a single flag to train on a GPU machine then deploy to commodity clusters or mobile devices. Speed makes Caffe perfect for research experiments and industry deployment.

### V. EXPERIMENTAL RESULTS AND DISCUSSION

The network structures used for classification are SVM and DT. In addition, Deep Learning (LeNet) was used. The results are shown in Table II. Here, ‘Medium Gaussian SVM’ is that type of a Gau<u>ss</u>ian-Kernel-SVM, for which, the kernel scale is set to the 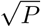, where *P* is the number of predictors.

**TABLE II.**
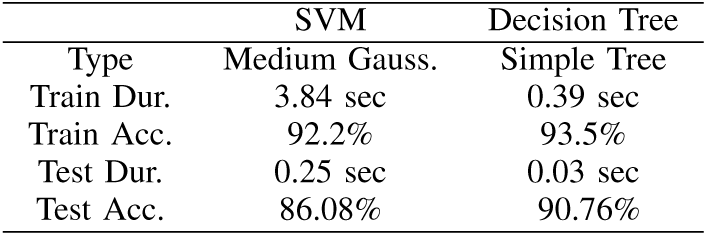
CLASSIFICATION RESULTS FOR 9×9 FEATURE IMAGES

LeNet deep learning model accepts 28×28 pixels image patches. In order to comply with the LeNet specification, the last row and the column of the 27×27 pixels feature image is zero-padded by one pixel. For each feature images (9 ×9 pixels and 27 ×27 pixels) Stochastic Gradient Descent (SGD) optimization is used in network. In this study, training and testing experiments are processed on a computer with Intel i7 6700 processor, 32 GB RAM, Asus Geforce GTX1080TI graphics card, and 1 TB HDD. The results are shown in Table III.

**TABLE III.** COMPARISON RESULTS FOR DEEP LEARNING – LENET

| | $9 \times 9$ feature image | $27 \times 27$ feature image |
| --- | --- | --- |
| Train Dur. | 23 sec | 23 sec |
| Train Acc. | 96.56% | 97.31% |
| Test Dur. | 6.4 sec | 6.4 sec |
| Test Acc. | 92.9% | 93.38% |

The size of the parameter space of the CNN structure is on the order of 105. However, due to the labelling burden, the current number of training samples are orders of magnitude less than the size of the parameter space. Currently, we are increasing the number of training samples. Despite the possibility of underfitting, the current standard LeNet structure yields acceptable classification results. Apart from increasing the number of training samples, our goal is to customize the network and have it comply with the myelin quantification application.

## VI. CONCLUSION

MS is a neurodegenerative disease without a cure. In order to develop a therapy for MS and other myelin related neurodegenerative diseases, evaluating effects of hundreds of thousands of chemicals on myelin is essential. Such an undertaking requires a rapid and efficient method to quantify myelin. Therefore, the aim of this work is to develop an automated myelin detection and quantification method.

Previously, myelin was segmented by identification of the overlapping regions of the neurons and oligodendrocytes [3]. These methods neither consider nor provide information on the shape of myelin. More, they may suffer from false positives as not all the overlaps are filtered out by the initial criteria set. In this study, we compared three different machine learning strategies to evaluate their effectiveness in segmenting myelin. The ground truths were extracted manually from images already classified by CEM. One hour of manual segmentation yields approximately 25 ground truths. Thus, extraction of a training set may take several day depending on the number of myelin contained in the image, reflecting the tremendous time and manpower required for manual quantification.

Next, both the ground truth and test images are converted into feature images as described above. In order to reflect the 3D nature of myelin, the 2D feature images were composed of three different z-sections. The multi-parameter of myelin recognition is considered by merging oligodendrocyte, neuron, and nucleus images. Finally, the images were classified using SVM, DT and CNN methods. The LeNet method yielded the most accurate results. Moreover, myelin classification using deep learning, the DeepMQ, took seconds compared to days of manual classification. In conclusion deep learning offers a novel avenue to explore for segmentation of biological structures that are composed of subcellular structures protruding from more than one cell type and whose defining criteria is the peculiar arrangement of these cellular protrusions relative to each other.

## ACKNOWLEDGMENT

We gratefully thank TUBITAK (project number: 316S026) and Turkish Academy of Sciences for their financial support.

